# New insights into functional divergence and adaptive evolution of uncultured bacteria in anammox community by complete genome-centric analysis

**DOI:** 10.1101/2023.08.15.553441

**Authors:** Yi-Cheng Wang, Yanping Mao, Hui-Min Fu, Jin Wang, Xun Weng, Zi-Hao Liu, Xiao-Wei Xu, Peng Yan, Fang Fang, Jin-Song Guo, Yu Shen, You-Peng Chen

## Abstract

Anaerobic ammonium-oxidation (anammox) bacteria play a crucial role in global nitrogen cycling and wastewater nitrogen removal, but they share symbiotic relationships with various other microorganisms. No pure culture is available for anammox bacteria so far. Although shotgun metagenomics based on short reads has been widely used in anammox research, metagenome-assembled genomes (MAGs) are often discontinuous and highly contaminated, which limits in-depth analyses of anammox communities. Here, for the first time, we performed Pacific Biosciences high-fidelity (HiFi) long-read sequencing on the anammox granule sludge sample from a lab-scale bioreactor, and obtained 30 accurate and complete metagenome-assembled genomes (cMAGs). These cMAGs were obtained by selecting high-quality circular contigs from initial assemblies of long reads generated by HiFi sequencing, eliminating the need for Illumina short reads, binning, and reassembly. One new anammox species and species from three novel families were found in this anammox community. cMAG-centric analysis revealed divergences in general and nitrogen metabolism among members of the anammox community. Furthermore, we identified mobile genetic elements (MGEs) and putative horizontal gene transfer (HGT) events within these cMAGs to explore the adaptive evolution of the community. The results suggest that MGEs and HGT events, particularly transposons containing *tnpA* in anammox bacteria, might play important roles in the adaptive evolution of this anammox community. The cMAGs generated in the present study could be used to establish of a comprehensive database for anammox bacteria and associated microorganisms. Our findings highlight the advantages of HiFi sequencing for the studies of complex mixed cultures such as anammox communities and advance our understanding of anammox communities.

## 1. Introduction

Anaerobic ammonium-oxidation (anammox) plays a key role in the global nitrogen cycle and is a cost-effective technique for treating ammonium-rich wastewaters due to no external carbon source requirement, lower oxygen consumption and sludge production (Peeters and van Niftrik, 2019; Adams et al., 2022). Approximately 120 anammox wastewater treatment plants are currently operating worldwide (Sheng et al., 2020). Anammox bacteria affiliated with the phylum Planctomycetota (Planctomycetes) are responsible for catalyzing anammox process, in which anammox bacteria utilize nitrite as electron acceptor to oxidize ammonium to dinitrogen gas (N_2_) under anoxic conditions (Kartal et al., 2013). Although anammox bacteria were discovered more than 20 years ago, and substantial efforts have been made to isolate them, yet pure cultures of these bacteria are still unavailable to date (Peeters and van Niftrik, 2019; Kuenen, 2020). It is well known that anammox bacteria hardly survive alone and share symbiotic relationships with various other microorganisms affiliated with the phyla Acidobacteriota, Armatimonadota, Bacteroidota, Chloroflexota, Proteobacteria, Patescibacteria, etc. (Bhattacharjee et al., 2017; Lawson et al., 2017; Feng et al., 2018; Zhao et al., 2018a). The co-existing bacteria can provide anammox bacteria with growth factors or stable nitrite under specific C/N ratios, and prevent the inhibition of anammox bacteria by organic matter (Zhao et al., 2018b; Cao et al., 2021). Therefore, comprehensive analysis of the diversity, functional divergence, and adaptive evolution of anammox communities can contribute to advance our understanding of anammox systems and promote the widespread application of this energy-efficient treatment process.

Shotgun metagenomics employing next-generation sequencing (NGS) has been widely used in studies of anammox systems, and the genomes of different anammox bacteria and associated bacteria have been successfully retrieved from samples from various habitats (Speth et al., 2016; Lawson et al., 2017; Suarez et al., 2022; Yang et al., 2022). However, due to the short reads and PCR requirement of NGS represented by the Illumina platform, leading to some shortcomings such as fragmented assembly and high genome contamination (Meziti et al., 2021). The recovered metagenome-assembled genomes (MAGs) generated by NGS are generally discontinuous because of repetitive, mobile, and conserved sequences, which limits their application in further analyses (Yuan et al., 2015; Sangwan et al., 2016; Meziti et al., 2019; Maguire et al., 2020). Long-read third-generation sequencing (TGS) developed by Pacific Biosciences (PacBio) and Oxford Nanopore Technologies (ONT) is a promising approach for addressing the above problems. The complete circular genomes of anammox bacteria from highly enriched cultures of *Candidatus* Kuenenia stuttgartiensis (> 87% of the total biomass) were previous recovered using only long reads generated by PacBio RS II sequencing (Frank et al., 2018; Ding and Adrian, 2020). Additionally, Liu et al. (2020) employed a combination of Illumina and ONT sequencing to develop an iterative hybrid assembly (IHA) method, which enabled the retrieval of eight circular genomes from one sample of a partial nitrification anammox reactor. However, nucleotide accuracy and assembly bubbles were not reported for these eight circular genomes. Consequently, a simpler and more efficient approach for recovering numerous accurate circular genomes from anammox systems is currently lacking. Furthermore, whether TGS can be employed alone to retrieve accurate and complete prokaryotic genomes, beyond those of dominant species, from anammox communities remains unclear.

Recently, PacBio high-fidelity (HiFi) sequencing, which can yield high-accuracy long reads (accuracy > 99.5% and average read length of 10–25 kb) (Hon et al., 2020), has become popular for the assembly of plant and animal genomes (Hon et al., 2020; Nurk et al., 2022), as well as the analysis of complex microbiomes (from human gut, sheep fecal, chicken cecum, and saline lake sediment samples) (Bickhart et al., 2022; Feng et al., 2022; Kim et al., 2022; Tao et al., 2023). HiFi sequencing has been shown to greatly improve the quality of metagenome assembly (Tao et al., 2023). Most importantly, HiFi sequencing enables the acquisition of accurate and complete prokaryotic genomes from complex microbiomes, without the need for Illumina short reads, binning, and reassembly. Kim et al. (2022) assembled the HiFi reads derived from five human gut samples, and obtained 102 complete metagenome-assembly genomes (cMAGs). One of these cMAGs is almost identical to the genome of pure culture, with an average nucleotide identity (ANI) value exceeding 99.99%. Taken together, HiFi sequencing provides a highly suitable approach for the accurate and comprehensive analysis of environmental microbiomes, such as anammox microbiomes. The utilization of HiFi sequencing enables the retrieval of more reliable cMAGs compared with MAGs. These cMAGs can serve as a valuable resource for subsequent downstream analyses, including the identification of novel species, exploration of genes associated with specific pathways (Parks et al., 2017; Galambos et al., 2019), investigation of functional divergence, and study of the adaptive evolutionary mechanisms of prokaryotes (Douglas and Langille, 2019).

To address the aforementioned research gaps and facilitate in-depth analysis of anammox communities, HiFi sequencing were performed on the granular sludge sample obtained from a lab-scale anammox bioreactor for the first time. The long-read metagenome obtained was *de novo* assembled, and non-redundant 30 cMAGs, including a genome of a novel anammox species and three genomes from novel families, were filtered out without any binning and reassembly processes. Using these reliable cMAGs, functional divergence and adaptive evolution within the anammox community were further explored. Overall, the results of this study provide a valuable workflow for analyzing complex aquatic microbiomes as well as new insights into bacterial interactions and evolution within the anammox community.

## 2. Materials and methods

### 2.1. Sample collection and DNA extraction

The anammox granular sludge samples were collected in quintuplicate from a lab-scale expanded granular sludge blanket (EGSB) reactor (**Fig. S1**), which has been operated for two years. The composition of the culture medium is shown in **Table S1** and **Table S2**. Residual sludge from a municipal wastewater treatment plant was used as the seed sludge. The EGSB reactor was operated at room temperature (the water bath portion was maintained at 35℃) and equipped with an effluent reflux system. Total genomic DNA was extracted using the E.Z.N.A.^®^ Soil DNA Kit (Omega Bio-Tek, USA). The purity and quantity of genomic DNA were assessed using the NanoDrop 2000 (ThermoFisher Scientific, USA) and the Qubit 3.0 Fluorometer (Invitrogen, USA) respectively (**Table S3**), and its integrity was verified using agarose gel electrophoresis (**Fig. S2**). The DNA sample with a high molecular weight and sufficient quantity was used in subsequent procedures.

### 2.2. Metagenomic sequencing

Metagenomic sequencing libraries were prepared and sequenced by Biozeron Biotechnology Co. Ltd. (Shanghai, China). For PacBio HiFi sequencing, 5 μg of DNA was used to construct a SMRTbell library with the SMRTbell Prep Kit (Pacific Biosciences, USA). Damaged double-stranded DNA in the initial DNA sample was repaired using the PreCR^®^ Repair Mix Kit (New England Biolabs, USA) before library construction. The repaired DNA was then size-selected to obtain molecules larger than 3 kb. The SMRTbell library was sequenced on the PacBio Sequel IIe platform using a single SMRT 8M cell. HiFi reads were generated with the CCS module of SMRT Link v11.0 (Pacific Biosciences, USA). For Illumina sequencing, the sequencing library was constructed with a fragment length of ∼450 bp and sequenced using the PE150 strategy on the Illumina NovaSeq 6000 platform. Basic information of both the HiFi and Illumina reads is shown in **Table S4** and **Table S5**, respectively.

### 2.3. *De novo* assembly and quality assessment

*De novo* assembly of HiFi reads was performed using three different assemblers: hifiasm-meta v0.3-r063.2 (Feng et al., 2022), Flye v2.9.1-b1780 (Kolmogorov et al., 2020), and Canu v2.2 (Nurk et al., 2020). Both the hifiasm-meta and Flye assembly were conducted using the recommended PacBio HiFi meta mode. The Canu assembly was conducted using the PacBio HiFi mode with recommended parameters. The assembly quality was assessed using QUAST v5.2.0 (Gurevich et al., 2013), and the results are shown in **Table S6**.

### 2.4. Filtering, taxonomic classification, and functional annotation of cMAGs

The HiFi cMAGs were filtered from initial assemblies using an adapted workflow published before (Kim et al., 2022). The workflow is briefly described as follows: (1) the circular contigs without assembly bubbles (assessed by BubbleGun v1.1.6 (Dabbaghie et al., 2022) as required) and repeats were filtered from the three assemblies; (2) contigs with sequence lengths shorter than 100 kb were removed; (3) the completeness and taxonomic classification of remaining contigs were assessed using GTDB-Tk v2.1.1 (Parks et al., 2022), and contigs with bacterial marker genes (bac120) less than 80 were removed (at this step, no archaea genome was found); (4) the SSU rRNA of the remaining contigs was predicted using barrnap v0.9 (https://github.com/tseemann/barrnap), and contigs missing 5S, 16S or 23S rRNA were removed; (5) the tRNA of remaining contigs was identified using tRNAscan-SE v2.0.11 (Chan et al., 2021), and contigs with tRNA types fewer than 18 were filtered out; and (6) ANI values of contigs were calculated using FastANI v1.33 (Jain et al., 2018). Redundant contigs were those with ANI values ≥ 99% and alignment percentages ≥ 95%, and the contigs with the maximal sum of ANI values were selected as the representative sequences. The contigs obtained through above six steps were considered non-redundant cMAGs. In addition, PlasClass v0.1 (Pellow et al., 2020) was used to determine whether the filtered circular contig represented a plasmid or not. The cMAG abundances represented by transcripts per million (TPM) were calculated using salmon v1.10.1 (Patro et al., 2017), by mapping Illumina reads onto these genomes.

Gene prediction of cMAGs was performed using Prodigal v2.6.3 (Hyatt et al., 2010), and the predicted genes were comprehensively annotated using MEBS v1.2 (De Anda et al., 2017), eggNOG database v5.0.2 (e-value < 1e-5) (Huerta-Cepas et al., 2019), NCycDB (e-value < 1e-5, identity > 40%, and coverage > 40%) (Tu et al., 2019), and KofamKOALA (Aramaki et al., 2020). The annotation results based on eggNOG database included the functional annotation of the Clusters of Orthologous Groups proteins (COGs) (Galperin et al., 2015) and CAZy (Lombard et al., 2014) databases. The gene abundances represented by TPM were also calculated using salmon v1.10.1 (Patro et al., 2017).

### 2.5. Identification of mobile genetic elements (MGEs) and horizontal gene transfer (HGT)

MGEs of cMAGs were identified by alignment (--escore = 1e-6, --pidentvalue = 80, and –queryscore = 75) against mobileOG-db beatrix-1.6 (Brown et al., 2022). The HGT events among all cMAGs were identified using MetaCHIP v1.10.12 (Song et al., 2019) with default parameters. All putative transferred genes were further annotated by alignment (e-value < 1e-5) against both the eggNOG database v5.0.2 (Huerta-Cepas et al., 2019) and NCBI non-redundant (NR) database.

### 2.6. Comparative genomic analyses

Reference genomes (**Table S7**) of 26 anammox bacteria in eight genera (*Ca.* Anammoxibacter, *Ca.* Anammoxoglobus, *Ca.* Anammoximicrobium, *Ca.* Brocadia, *Ca.* Jettenia, *Ca.* Kuenenia, *Ca.* Loosdrechtia, and *Ca.* Scalindua) were downloaded from both the NCBI and EMBL-EBI databases. JSpeciesWS (Richter et al., 2015) was used to calculate the ANI values (based on blast) between cMAGs representing anammox bacteria and other reference genomes of *Ca.* Jettenia. DNA-DNA hybridization (DDH) *in silico* was performed using GGDC v3.0 (Meier-Kolthoff et al., 2022) with recommended parameters. All reference anammox bacterial genomes were also comprehensively annotated like cMAGs.

## 3. Results

### 3.1. Overview of cMAGs

A total of 541 circular contigs were obtained through HiFi sequencing and assembly (**Fig. 1)**. These contigs were filtered based on biological features and ANI values, yielding 30 non-redundant contigs known as cMAGs (**Fig. 1)**. As shown in **Table S8**, the predicted values by PlasClass for all cMAGs were far lower than the threshold of 0.5, thus none of them represented plasmids. The number of cMAGs from the assembly of hifiasm-meta, Canu and Flye was 19, 7, and 4, respectively. The size of these cMAGs ranged from 0.6 to 4.4 Mb, and the number of coding sequences (CDSs) ranged from 629 to 4,388 (**Table 1**). According to the taxonomic classification by GTDB-Tk, only 16 cMAGs were assigned to known species (**Fig. 2**). Three cMAGs could not be assigned to known families, and four cMAGs could not be assigned to known genera. These cMAGs represented nine phyla, with Patescibacteria having the highest number of cMAGs. Notably, more than 75% and 50% of Patescibacteria genomes were not assigned at the species and genus level, respectively. The genomes of Patescibacteria bacteria were very small, with far fewer CDSs than the genomes of bacteria affiliated with other phyla (**Table 1**). The phyla Actinobacteriota, Armatimonadota, Bdellovibrionota, and Proteobacteria each had only one member, but the abundances of cMAG.1 (Actinobacteriota) and cMAG.30 (Proteobacteria) were not low compared with other cMAGs. cMAG.27, cMAG.1, and cMAG.25 were the most abundant in the community of 30 members, and the abundances of nine species (cMAG.10, cMAG.17, cMAG.5, cMAG.26, cMAG.18, cMAG.19, cMAG.2, cMAG.23, and cMAG.13) were less than one-fiftieth of that of cMAG.27. No pure cultures are available for any of the 16 species assigned at the species level at present, indicating that all cMAGs might represent uncultured species.

**Fig. 1.**
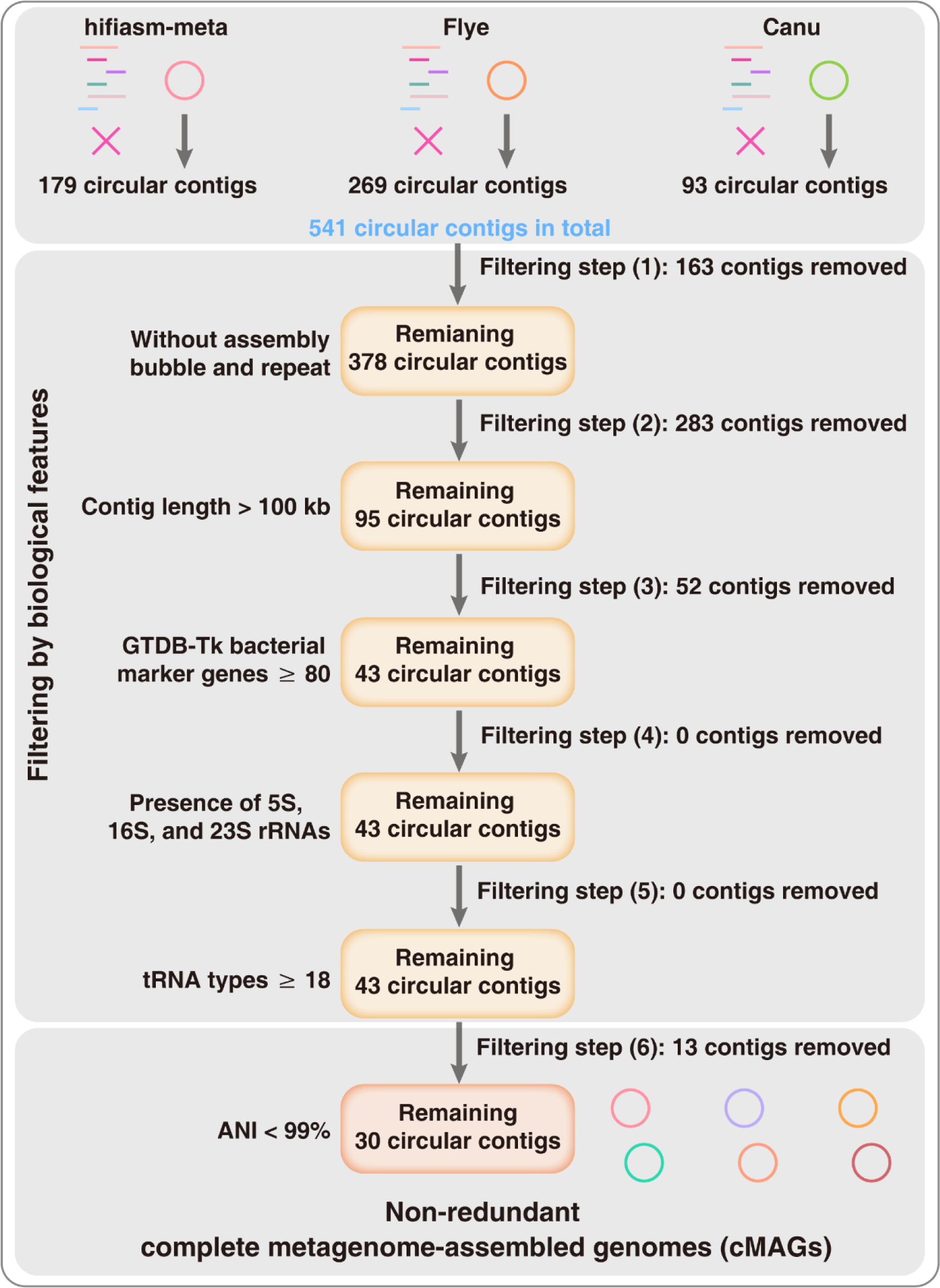
Workflow for filtering the HiFi cMAGs from three assemblies by hifiasm-meta, Flye, and Canu.

**Fig. 2.**
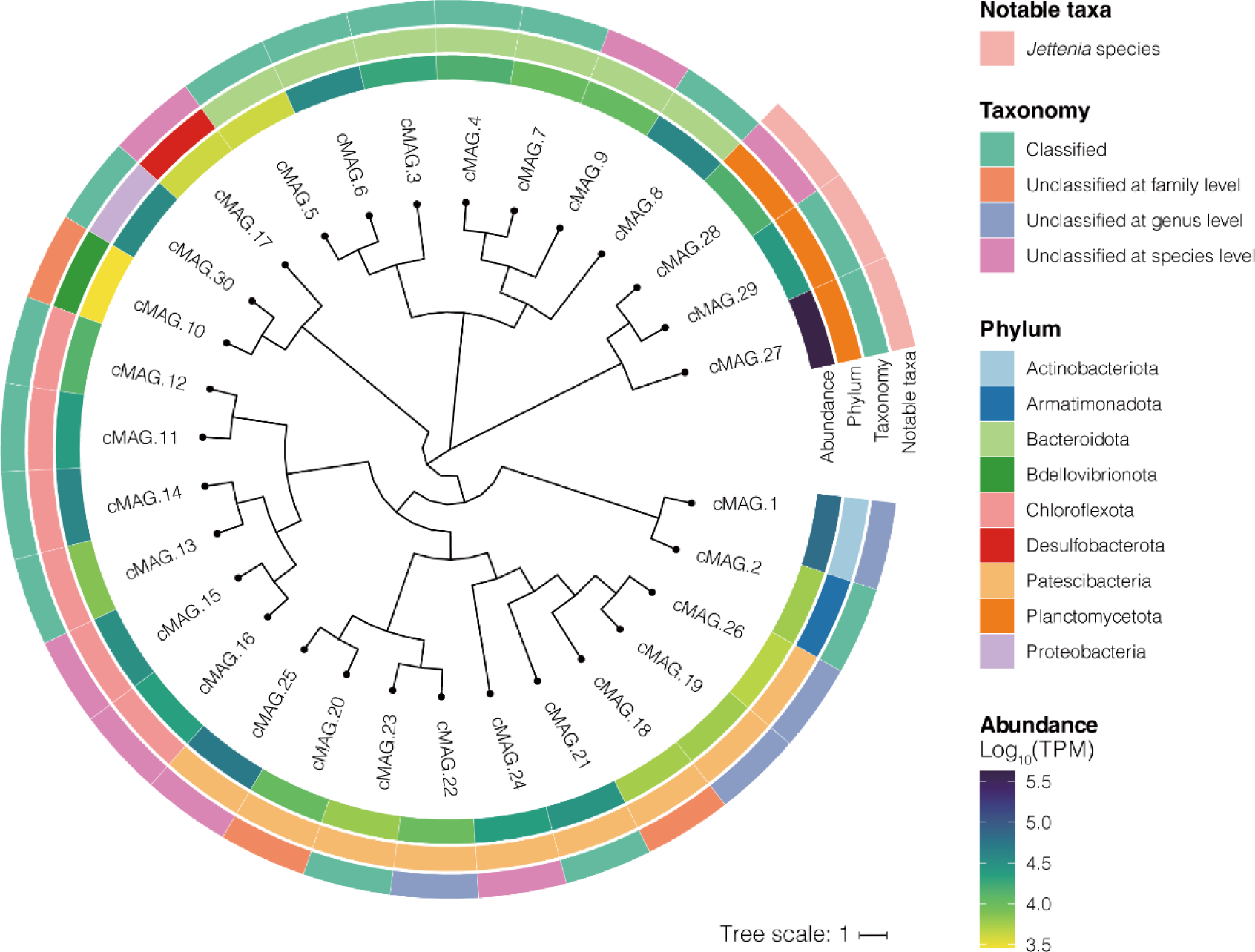
The phylogenetic placement and abundance of cMAGs recovered from anammox granular sludge. The phylogenetic tree was inferred based on the multiple sequence alignments of conserved marker genes from 30 cMAGs using the GTDB-Tk.

**Table 1.** Characteristics of the 30 cMAGs obtained in this study.

| ID | Phylum (Genus) | Genome size (Mb) | GC (%) | CDS | GTDB-Tk marker gene | Types of tRNA | Features (5S/16S/23S) |
| --- | --- | --- | --- | --- | --- | --- | --- |
| cMAG.1 | Actinobacteriota (Unclassified) | 2.2 | 68.58 | 2,260 | 118 | 23 | 1/1/1 |
| cMAG.2 | Armatimonadota (H1-ARM1) | 2.9 | 60.95 | 2,651 | 111 | 23 | 1/1/1 |
| cMAG.3 | Bacteroidota (OLB6) | 3.0 | 49.64 | 2,427 | 119 | 22 | 2/2/2 |
| cMAG.4 | Bacteroidota (BCD5) | 3.3 | 36.15 | 2,825 | 119 | 22 | 1/1/1 |
| cMAG.5 | Bacteroidota (UTCHB3) | 3.4 | 42.13 | 2,623 | 118 | 23 | 1/1/1 |
| cMAG.6 | Bacteroidota (SURF-28) | 3.7 | 34.47 | 3,351 | 118 | 22 | 1/1/1 |
| cMAG.7 | Bacteroidota (OLB8) | 3.9 | 65.21 | 3,252 | 119 | 22 | 1/1/1 |
| cMAG.8 | Bacteroidota (PHOS-HE28) | 4.0 | 41.08 | 3,258 | 119 | 22 | 2/2/2 |
| cMAG.9 | Bacteroidota (UBA2336) | 4.1 | 43.59 | 3,752 | 117 | 23 | 2/2/2 |
| cMAG.10 | Bdellovibrionota (Unclassified) | 2.3 | 55.04 | 2,089 | 93 | 18 | 1/1/1 |
| cMAG.11 | Chloroflexota (UBA7227) | 3.5 | 59.60 | 3,200 | 116 | 24 | 1/1/1 |
| cMAG.12 | Chloroflexota (UBA7227) | 3.6 | 59.76 | 3,268 | 116 | 24 | 1/1/1 |
| cMAG.13 | Chloroflexota (UBA12294) | 3.9 | 53.51 | 3,590 | 111 | 23 | 1/1/1 |
| cMAG.14 | Chloroflexota (UBA12294) | 4.0 | 53.65 | 3,738 | 110 | 23 | 1/1/1 |
| cMAG.15 | Chloroflexota (OLB14) | 4.1 | 54.13 | 3,820 | 113 | 23 | 1/1/1 |
| cMAG.16 | Chloroflexota (OLB14) | 4.4 | 53.56 | 4,388 | 115 | 24 | 1/1/1 |
| cMAG.17 | Desulfobacterota (UBA5799) | 2.8 | 57.87 | 2,559 | 119 | 23 | 1/1/1 |
| cMAG.18 | Patescibacteria (Unclassified) | 1.0 | 44.78 | 956 | 89 | 20 | 1/1/1 |
| cMAG.19 | Patescibacteria (Unclassified) | 1.1 | 39.76 | 982 | 84 | 21 | 1/1/1 |
| cMAG.20 | Patescibacteria (Unclassified) | 1.2 | 60.26 | 1,204 | 95 | 21 | 1/1/1 |
| cMAG.21 | Patescibacteria (OLB21) | 1.2 | 38.42 | 1,073 | 93 | 21 | 1/1/1 |
| cMAG.22 | Patescibacteria (Unclassified) | 0.6 | 52.17 | 629 | 82 | 21 | 1/1/1 |
| cMAG.23 | Patescibacteria (BOKC01) | 0.7 | 40.37 | 698 | 84 | 22 | 1/1/2 |
| cMAG.24 | Patescibacteria (UBA12026) | 0.8 | 34.42 | 947 | 80 | 20 | 1/1/1 |
| cMAG.25 | Patescibacteria (UBA919) | 0.9 | 36.84 | 827 | 90 | 22 | 1/2/1 |
| cMAG.26 | Patescibacteria (Unclassified) | 0.9 | 41.50 | 858 | 87 | 21 | 1/1/1 |
| cMAG.27 | Planctomycetota (*Ca. Jettenia*) | 3.5 | 41.60 | 3,145 | 108 | 23 | 2/2/2 |
| cMAG.28 | Planctomycetota (*Ca. Jettenia*) | 3.8 | 39.42 | 3,292 | 107 | 24 | 2/2/2 |
| cMAG.29 | Planctomycetota (*Ca. Jettenia*) | 4.0 | 40.01 | 3,510 | 107 | 24 | 2/2/2 |
| cMAG.30 | Proteobacteria (QUBU01) | 3.4 | 66.68 | 3,021 | 118 | 23 | 1/1/1 |

Three anammox bacteria, namely cMAG.27, cMAG.28, and cMAG.29, were identified and they were all assigned to the genus *Ca.* Jettenia. According to the classification by GTDB-Tk, cMAG.28 was unclassified at the species level, and cMAG.27 and cMAG.29 were assigned to the species *Ca.* Jettenia sp. AMX2 and *Ca.* Jettenia caeni, respectively. Therefore, we downloaded the 11 available *Ca.* Jettenia genomes from the NCBI database and calculated ANI values among the above three cMAGs and other *Ca.* Jettenia genomes. As shown in **Fig. S3**, ANI values between cMAG.29 and *Ca.* Jettenia caeni (GCA_000296795.1), and between cMAG.27 and *Ca.* Jettenia sp. AMX2 (GCA_900696655.1) were 99.70% and 99.45%, respectively. These results indicate that cMAG.29 and cMAG.27 are same species of *Ca.* Jettenia caeni and *Ca.* Jettenia sp. AMX2, respectively. The ANI value between cMAG.28 and *Ca.* Jettenia sp. AM49 (GCA_021650895.1) was 95.83%, which was just within the threshold of 95–96% species boundary (Richter and Rosselló-Móra, 2009; Palmer et al., 2020). Consequently, we calculated the DDH value between cMAG.28 and *Ca.* Jettenia sp. AM49 (GCA_021650895.1), and the resulting DDH value (66.60%) was less than the species boundary threshold of 70% (Meier-Kolthoff et al., 2013), indicating that cMAG.28 could represent a novel species of *Ca.* Jettenia different from *Ca.* Jettenia sp. AM49.

### 3.2. Functional divergence in general metabolism

Metabolic inference revealed that, aside from members of Patescibacteria, other bacteria in the anammox system might play a crucial role in nitrogen (N) and iron (Fe) cycles (**Fig. 3**). Additionally, members of phyla such as Planctomycetota and Chloroflexota were found to be important for the carbon (C) cycle in this anammox system. These inferences were further supported by the presence of important metabolic genes or pathways in their genome. For instance, members of the phyla Planctomycetota and Chloroflexota harbor a relatively high abundance of carbohydrate enzymes, including glycoside hydrolases and glycosyl transferases (**Fig. S4**).

**Fig. 3.**
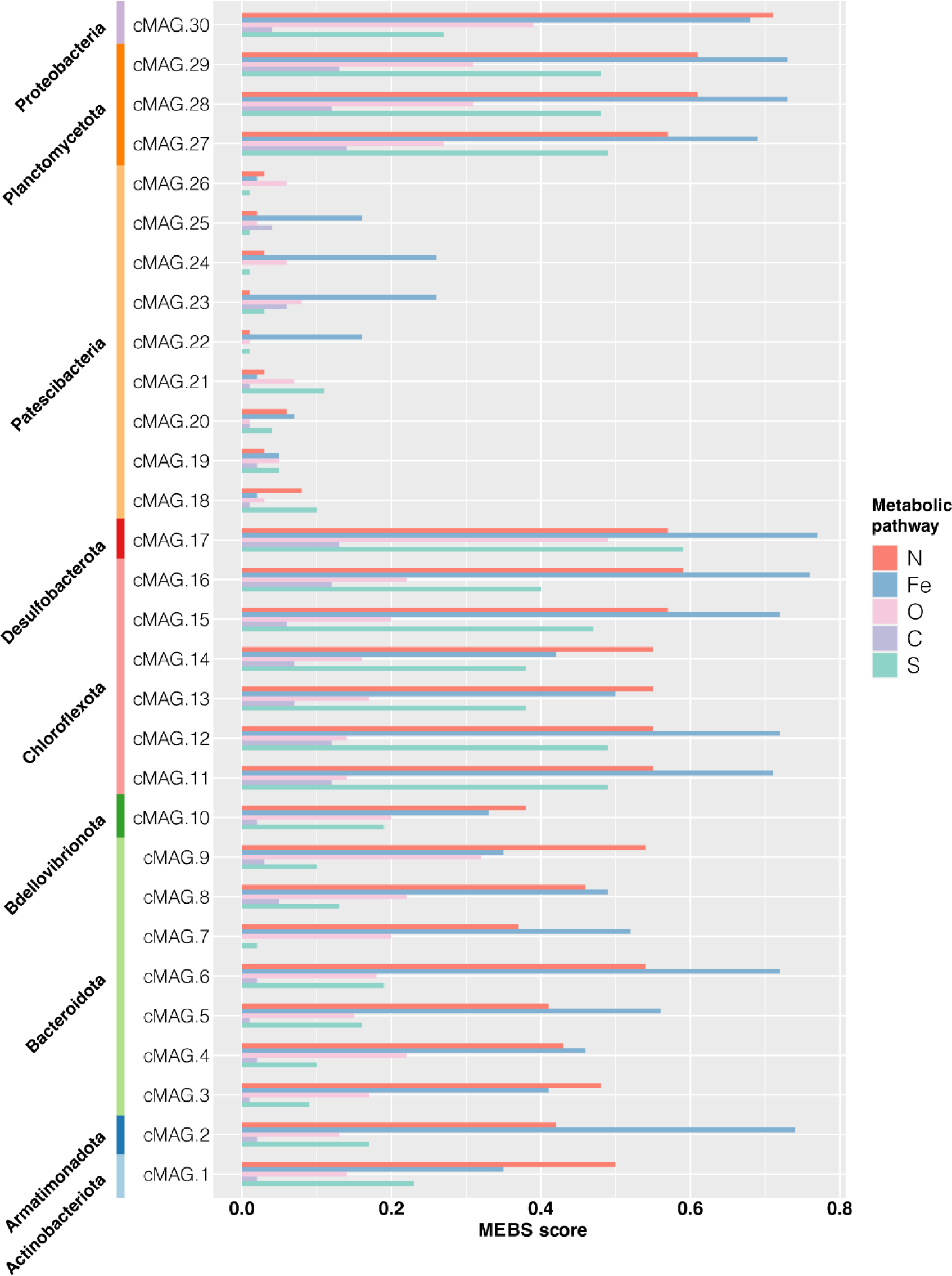
The MEBS scores for the nitrogen (N), iron (Fe), oxygen (O), carbon (C) and sulfur (S) metabolic pathways of 30 cMAGs.

In this community, there might be 13 species capable of autotrophic growth that utilize three different carbon fixation pathways. Conversely, no known carbon fixation pathways could be identified in the remaining 17 species, suggesting that they might be heterotrophic. In detail, all the members of the phyla Chloroflexota and Planctomycetota (cMAG.27, cMAG.28, and cMAG.29) harbor the key genes encoding CO dehydrogenase and acetyl-CoA synthase, which are essential for carbon fixation via the Wood-Ljungdahl (WL) pathway. As for the only Proteobacteria bacterium (cMAG.30), the presence of a nearly complete Arnon-Buchanan cycle indicates its potential for autotrophic growth. Moreover, the presence of the complete Embden– Meyerhof pathway (EMP) and the non-oxidative pentose phosphate pathway (PPP) pathways in cMAG.30 reflect its capability for heterotrophic growth. Besides, the presence of genes encoding cytochrome c oxidase (cbb_3_-type), cytochrome bc_1_ ubiquinol oxidase, and succinate dehydrogenase in cMAG.30 suggests its capability for aerobic respiration. Three members of Patescibacteria (cMAG.18, cMAG.19, and cMAG.26) were found to carry the RubisCO-encoding gene (*rbcL*), suggesting that they might perform carbon fixation through the Calvin-Benson-Bassham cycle (CBB). The presence of complete EMP and the non-oxidative PPP pathways in three *Ca.* Jettenia species (cMAG.27, cMAG.28, and cMAG.29), as well as a complete tricarboxylic acid cycle (TCA cycle) and *pdhA* encoding pyruvate dehydrogenase, suggest their capabilities for fermentation.

### 3.3. Functional divergence in nitrogen metabolism

To further explore functional divergence in nitrogen metabolism of these bacteria, 30 cMAGs were comprehensively annotated using a curated nitrogen cycle database (NCycDB). A total of 24 species were identified as potentially participating in key nitrogen conversion processes, such as anammox, denitrification, nitrification, and dissimilatory nitrate reduction to ammonia (DNRA) (**Fig. 4A and B**). Three members of *Ca.* Jettenia harbor the full set of anammox genes, including *nirK*/*nirS*, *hzs*, and *hdh*, but they all lack *nxrA* encoding nitrite oxidoreductase (**Fig. 4A**). In addition, they all harbor *napA, narG*, and *nrfA*, suggesting their capacities for DNRA. Similarly, among the 11 *Ca.* Jettenia species from other habitats, none of them carry the *nxrA* gene (**Fig. S5**). Only three members, i.e. two members of *Ca.* Jettenia caeni and one member of *Ca.* Jettenia sp. AMX2, harbor both *napA* and *narG*. Nearly all of these *Ca.* Jettenia members (10 out of 11) may be capable of DNRA. Members belonging to the genera *Ca.* Jettenia and *Ca.* Brocadia possess a richer repertoire of nitrogen metabolism marker genes compared with anammox bacteria in other genera (**Fig. S5**).

**Fig. 4.**
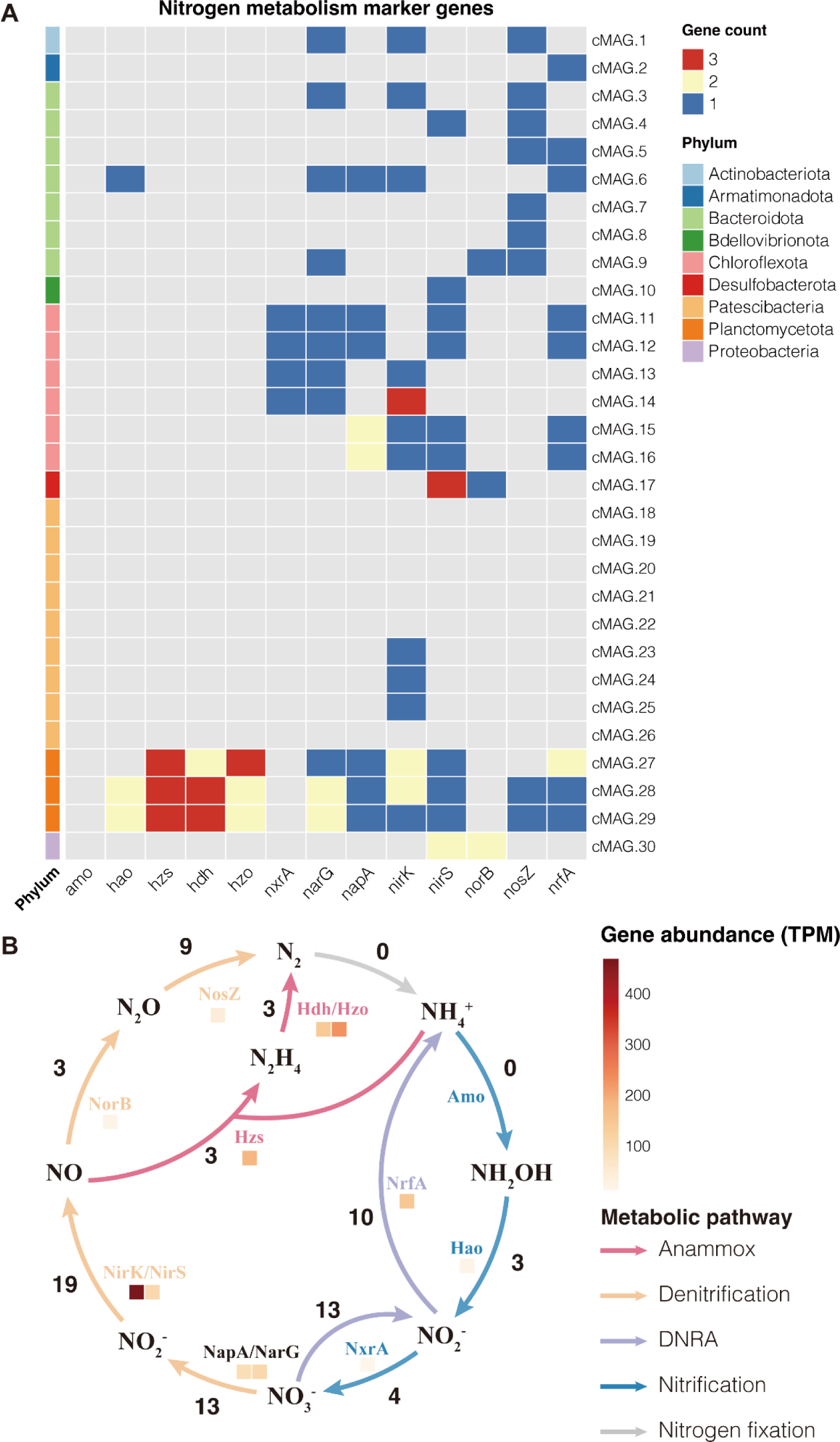
The potential for nitrogen metabolism of 30 cMAGs. (**A**) Nitrogen metabolism marker genes. (**B**) Proposed pathway for nitrogen conversion in the anammox community. The number indicates the number of species with genes encoding a given step. Colored squares indicate gene abundances.

A total of 20 non-anammox bacteria community members were found to carry denitrification genes (**Fig. 4A**). However, none of them carry the full set of denitrification genes, indicating a potential for cooperative denitrification. In addition, seven members carry the key gene *nrfA* for DNRA, while only five of them harbor the complete DNRA pathway. The Armatimonadota bacterium (cMAG.2) exclusively harbors *nrfA*. According to gene abundance, the denitrification pathway might have not outcompeted both the anammox and DNRA pathways (**Fig. 4B)**. Of note, the majority of Patescibacteria members were not been identified with important known nitrogen metabolism genes, suggesting that they might play other roles in the system. No complete nitrification pathway was identified in any of these cMAGs, and the gene encoding ammonia monooxygenase (*amo*) was not found.

### 3.4. Metagenomic evidence for adaptive evolution

To gain insights into the adaptive evolution of the anammox community, an analysis was conducted to identify MGEs in all cMAGs using the latest MGE database (mobileOG-db) with stringent parameters. Among these cMAGs, eight members were found to carry MGEs, with a total of 63 MGEs identified (**Fig. 5A**). Notably, cMAG.4, cMAG.7, cMAG.28, and cMAG.30 were found to harbor sequences related to phages. The majority of the predicted MGEs were associated with integration or excision processes. Of note, one of the anammox bacteria (cMAG.29) harbored a discrete distribution of 28 transposons, all of which contained the transposase-encoding gene *tnpA* (**Fig. S6**). Another anammox bacterium (cMAG.28) also harbors three transposons that containing *tnpA*, while the remaining members (including cMAG.27) did not carry *tnpA*. The same identification procedure was performed on the other 26 reference genomes of anammox bacteria. Approximately half of the anammox bacteria carried *tnpA*, with gene counts no less than 18 in *Ca.* Jettenia caeni (only one) and all members of *Ca.* Kuenenia stuttgartiensis, and others carried no more than eight *tnpA* genes (**Fig. 5B**). Furthermore, the upstream and downstream genes of *tnpA* in cMAG.29 were found to be different (**Fig. 6**), indicating that they represent the simple transposon (insertion sequence) without additional functional genes.

**Fig. 5.**
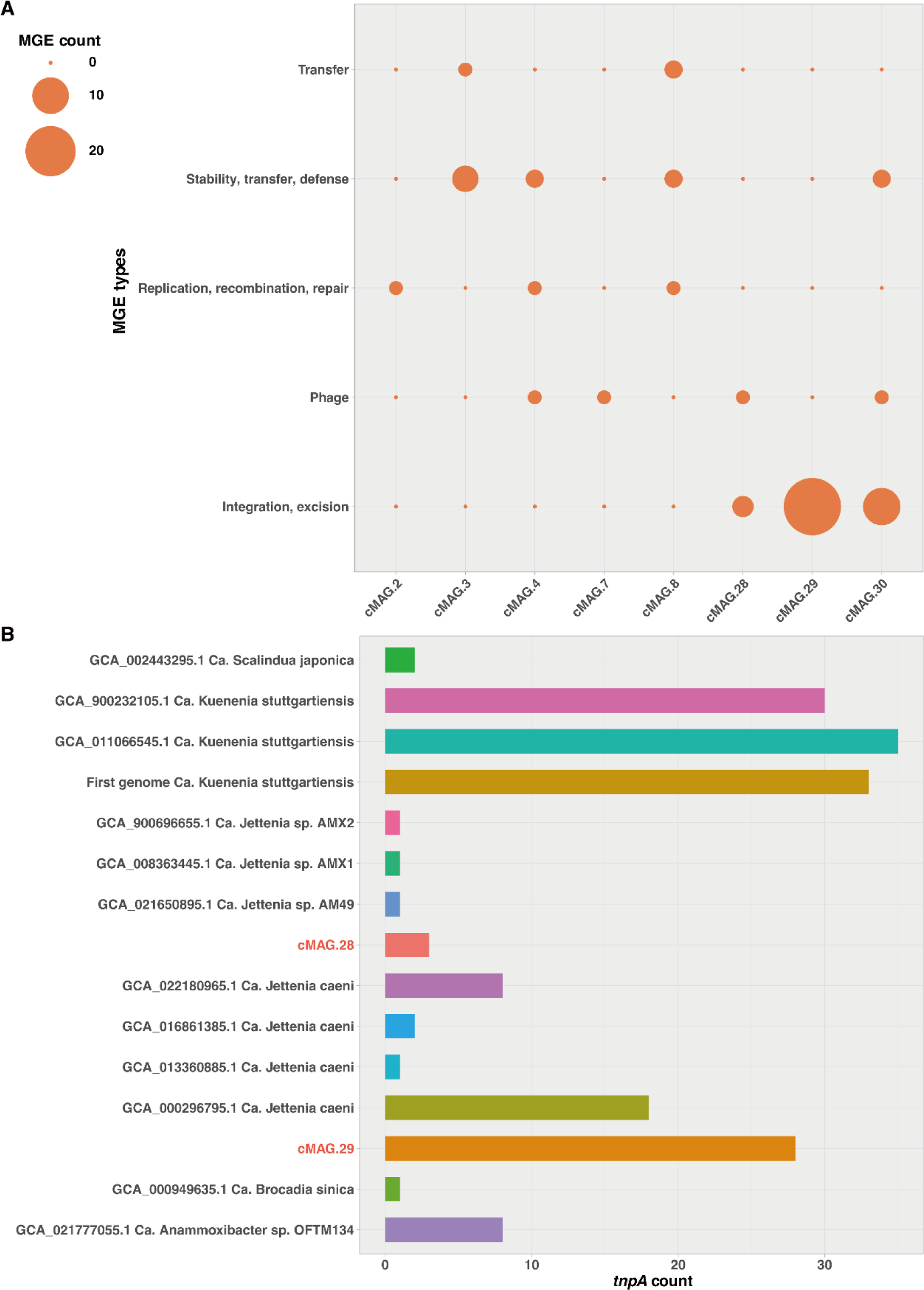
Mobile genetic elements (MGEs) identified in the cMAGs and reference genomes. (**A**) MGEs identified in cMAGs. (**B**) The *tnpA* counts identified in two *Ca.* Jettenia species of this study and other anammox species.

**Fig. 6.**
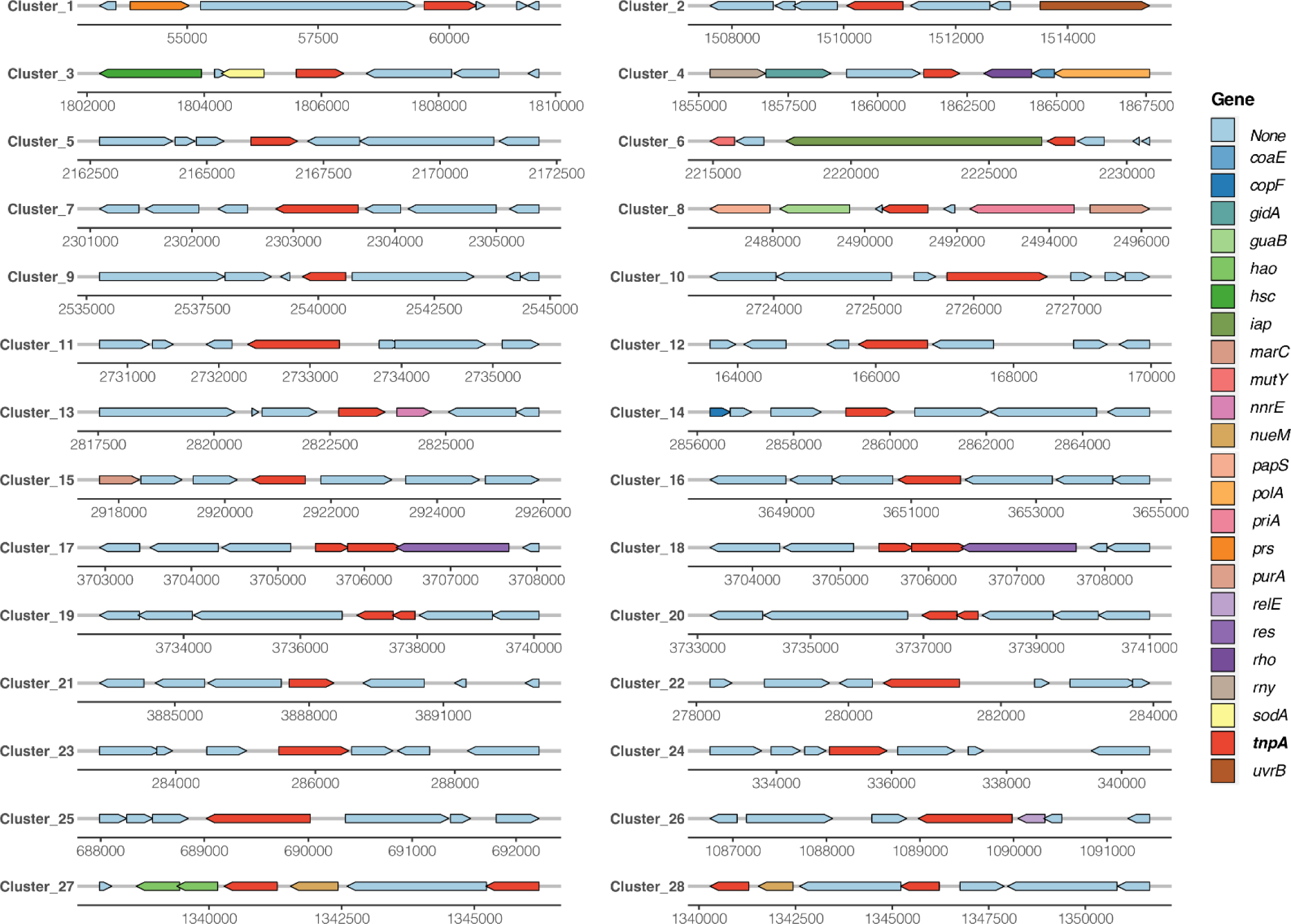
The predicted 28 gene clusters containing *tnpA* of cMAG.29.

In addition to MGE analysis, HGT events were identified on all cMAGs. A total of 50 putative HGT events along with their transfer direction were identified, involving 6 phyla and 14 species (**Fig. 7A**). Most of the putative HGT events occurred within the phylum Chloroflexota, accounting for 66% of the total. Putative HGT events within the phylum Bacteroidetes accounted for 34% of the total, with 24% assumed to occur with members of different phyla. At the species level, putative HGT events were the most abundant in one member of the genus *UBA12294* (cMAG.13), with 25 donors and 5 recipients. Interestingly, the identification by MetaCHIP showed that the denitrifying bacterium cMAG.13 donated 10 genes to another denitrifying bacterium, cMAG.16, and reciprocally, cMAG.16 donated 5 genes to cMAG.13. A putative HGT event was observed between an anammox bacterium (cMAG.27) and a denitrifying bacterium affiliated with Bacteroidota, showing that cMAG.27 might acquire a gene from cMAG.8.

**Fig. 7.**
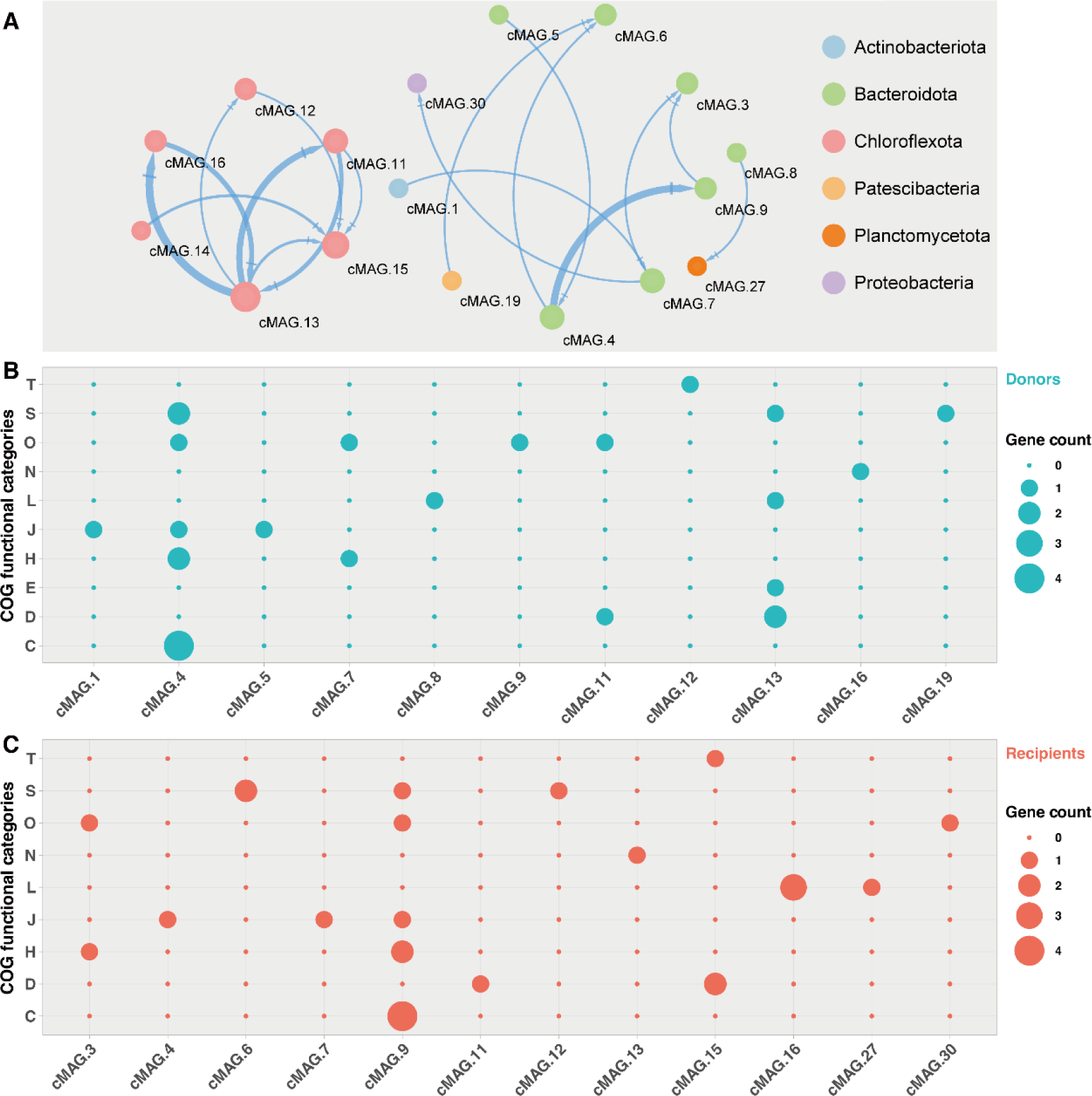
Network and functions of HGT genes of the anammox community. (**A**) HGT network of the anammox community; the direction and width of blue arrows indicate the direction of transfer and gene counts, respectively; the size of the node represents the count of connected edges. (**B**) and (**C**) Bubble size indicates the count of HGT genes annotated with COG functional categories. C: Energy production and conversion; D: Cell cycle control, cell division, chromosome partitioning; E: Amino acid transport and metabolism; H: Coenzyme transport and metabolism; J: Translation, ribosomal structure and biogenesis; L: Replication, recombination and repair; N: Cell motility; O: Posttranslational modification, protein turnover; S: Function unknown; T: Signal transduction mechanisms.

The donor and recipient genes for putative HGT events were identified. A total of 26 genes (> 50% of the total) in both donated and received genes could be annotated using the COG database (**Fig. 7B and C**). The donated genes are primarily responsible for energy production and conversion (C, 4 genes), posttranslational modification, protein turnover, chaperones (O, 4 genes), while four genes were annotated with unknown functions. For example, the Bacteroidota bacterium (genus *UBA2336*) cMAG.9 might acquire four genes (*sucD*, *fumC*, *atpA*, and *atpD*) from another Bacteroidota bacterium cMAG.4. It should be noted that the anammox bacterium cMAG.27 might acquire a gene encoding IS200 family transposase. No transfer genes directly related to nitrogen removal were identified. However, nearly half of the genes in both donors and recipients were either unannotated or had unknown functions; thus, additional investigation is required.

## 4. Discussion

In this study, a total of 30 complete circular prokaryotic genomes were successfully recovered from an anammox community using HiFi sequencing. We made slight adjustments to the filtering criteria compared with a previous study (Kim et al., 2022) to include circular contigs that did not meet all the proposed criteria but exhibited other desirable features. Since the accuracy of HiFi sequencing was extremely high (average quality = Q29.5) and the circular contig with assembly errors had been removed, the substandard circular contigs represented either chromosomes or circular plasmids. According to the prediction by PlasClass, these substandard contigs were not circular plasmid sequences. More importantly, most of these contigs were assigned to Patescibacteria, which are also referred to as candidate phyla radiation bacteria and are known to have much smaller genomes compared with other prokaryotes (Luef et al., 2015; Castelle and Banfield, 2018; Tian et al., 2020). They might have fewer marker genes than other conventional bacteria. These substandard contigs were therefore retained for downstream analysis. In strict terms, some of the recovered cMAGs from this study or previous studies (Liu et al., 2020; Kim et al., 2022) might represent complete chromosomes rather than complete genomes, as the existence of associated MGEs (e.g., plasmids) remains uncertain. For complex mixed cultures, such as anammox communities, accurately resolving the link of MGEs to their host complete chromosomes poses a major challenge. To address this problem, we propose that a combination of the recently developed high-throughput single-microbe sequencing (Microbe-seq) (Zheng et al., 2022) with HiFi sequencing would offer an effective strategy.

Although hifiasm-meta did not generate the highest number of circular contigs, it yielded cMAGs that accounted for 63% of the total. This is consistent with previous findings and highlights the advantages of hifiasm-meta for the assembly of HiFi reads (Kim et al., 2022). These cMAGs represent anammox bacteria and associated bacteria, including those with low abundance, and ∼47% of them may represent novel species, including a novel species of anammox bacteria. This suggests that a considerable number of anammox-associated bacteria remain poorly characterized. The reliability of the recovered cMAGs was supported by the high ANI values with conspecific genomes from previous studies. Our approach outperformed the IHA method (Liu et al., 2020) in terms of using fewer long reads (26.7 Gb vs. 69.4 Gb) and no need for short reads while ensuring the accuracy of circular genome reconstruction, demonstrating the potential for HiFi sequencing to accurately recover prokaryotic genomes from complex environmental microbiomes.

cMAGs provide comprehensive and accurate information on the metabolic potential and genetic interactions within the anammox community. Unlike previous MAGs (Lawson et al., 2017; Liu et al., 2020; Kallistova et al., 2022) that were affected by discontinuity and contamination, these cMAGs offer a more complete picture of the community. The anammox system was thought to be autotrophic due to the absence of external organic carbon source, but there were still many heterotrophs in this system. Actually, it is very common that anammox bacteria share symbiotic relationships with heterotrophic bacteria (Kindaichi et al., 2012; Lawson et al., 2017; Zhao et al., 2018b). In light of our culture conditions (without external organic carbon source), these co-existing heterotrophic bacteria might be responsible for granular aggregation, supplying growth factors to anammox bacteria, and consuming organic matter.

Approximately 70% of the species in the community were potentially engaged in nitrogen removal processes, either through anammox or denitrification pathways. The remaining 30% might not be directly involved in such nitrogen conversion processes. Nevertheless, some of them could perform carbon fixation, which provides organic matter to other co-existing bacteria. These members of Patescibacteria might also play roles in the degradation of poly-N-acetylglucosamine produced by anammox bacteria (Hosokawa et al., 2021). Three anammox bacteria affiliated with *Ca.* Jettenia could conduct DNRA and carbon fixation through the WL pathway, which is consistent with the results of previous studies (Strous et al., 2006; Lawson et al., 2017; Mardanov et al., 2019). Intriguingly, our results indicate that these three anammox bacteria were capable of respiratory ammonification with oxidation acetate, and similar results were also observed for the two other anammox bacteria affiliated with *Ca.* Jettenia (Ali et al., 2015; Mardanov et al., 2019). The DNRA capability of anammox bacteria could be a valuable strategy for survival, as they would no longer dependent on other microorganisms for the supply of ammonium and nitrite (Kartal et al., 2007). Furthermore, not all anammox bacteria are capable of carrying out the DNRA pathway, and it could be speculated that these anammox bacteria with the DNRA pathway might be better adapted to environments with a low concentration of ammonium and a high concentration of nitrite or nitrate.

Adaptive evolution plays a crucial role in the adaptation of bacteria to different habitats, and both MGEs and HGT contribute to this process (Brito, 2021; Arnold et al., 2022; Shi et al., 2022). We identified abundant MGEs and putative HGT events within the anammox community. Certain members of the community showed evidence of past phage infections, which led to mutations in their genomes. It is well known that transposons can provide a pool of pre-evolved genes that help bacteria adapt quickly to their environment (Dahlberg and Hermansson, 1995). The transposons containing the *tnpA* gene were prevalent in some anammox bacteria, indicating they might play important roles in their environmental adaptation. As mentioned above, anammox bacteria are not available for pure culture to date; thus, molecular biology methods based on monoclones cannot be used for the identification of metabolic pathways or bioaugmentation. Many questions about the basic metabolic pathways of anammox bacteria have not yet been addressed (Peeters and van Niftrik, 2019; Kuenen, 2020), challenges (such as the fast start-up and operation under non-optimal conditions) during the full-scale operation of anammox also require attention (Adams et al., 2020; Kouba et al., 2022), and this will require the development of methods for the genome editing of anammox bacteria. We believe that the transposon containing *tnpA* might be one of the breakthroughs for the genome editing of anammox bacteria in mixed cultures.

A considerable number of putative HGT events were predicted, suggesting that HGT might be another important strategy for promoting the adaptive evolution of bacteria within the anammox community. The transferred genes are involved in multiple metabolic pathways rather than antibiotic resistance, showing that these putative HGT events might contribute to the metabolic specialization and competitive advantage of these species (Seeleuthner et al., 2018; Wang et al., 2022). Particularly, the putative HGT events mainly occurred in denitrifying bacteria in the phyla Chloroflexota and Bacteroidota. Similar findings were also observed in the previous study on a full-scale anammox system (Wang et al., 2022). It could be inferred that denitrifying bacteria might be very active in genetic interactions within anammox communities. The putative HGT event between cMAG.27 and cMAG.8 shows that HGT events could occur between anammox bacteria and co-existing bacteria affiliated with distantly related lineages. Moreover, the transferred gene is a transposase-coding gene, again suggesting that some specific transposons could be utilized for the genome editing of anammox bacteria.

## 5. Conclusions

Our study demonstrates that HiFi sequencing enables the reconstruction of accurate cMAGs from the complex anammox community without the need for Illumina short reads, binning, and reassembly. These cMAGs represent not only high-abundance species, but also low-abundance species in this anammox community. Additionally, these cMAGs provide valuable insights into microbial diversity, metabolic divergence, and genetic interactions within the community, and they may lay a core foundation for a comprehensive database for anammox bacteria and associated microorganisms. The presence of MGEs and putative HGT events suggests ongoing evolution within the community, and they might play an important role in the domestication of the anammox culture. To our knowledge, this is the first study to describe the functional divergence and adaptive evolution of uncultured bacteria in an anammox community at the complete genome level. Further studies are warranted to confirm the functional divergence and genetic interactions within anammox communities, employing a combination of HiFi sequencing and other techniques.

## Data availability

The 30 cMAGs generated in this study have been deposited in the GenBank under the accession number PRJNA983097. The raw sequencing data have been deposited in the Sequence Read Archive under the accession number PRJNA981667 and PRJNA981689.

## Acknowledgments

This study was supported by the National Natural Science Foundation of China (21876016, 52070031, and 42207022) and the Natural Science Foundation of Guangdong Province (2023A1515012019).

