## Supplemental Table 1-8; Supplemental Figure 1-6 for "New insights into functional divergence and adaptive evolution of uncultured bacteria in anammox community by complete genome-centric analysis"

The supplementary information includes:

**Tables**

- **Table S1.** Composition of the culture medium.
- **Table S2.** Composition of trace element solution.
- **Table S3.** Assessment of the purity and quantity of genomic DNA.
- **Table S4.** Basic information of HiFi reads.
- **Table S5.** Basic information of Illumina reads.
- **Table S6.** The quality assessment of HiFi reads assemblies.
- **Table S7.** Reference genomes of 26 anammox bacteria.
- **Table S8.** The prediction results by PlasClass.

**Figures**

- **Fig. S1.** The lab-scale expanded granular sludge blanket (EGSB) reactor.
- **Fig. S2.** Agarose gel electrophoresis of qualified genomic DNA.
- **Fig. S3.** The average nucleotide identity (ANI) value heatmap among 3 cMAGs and other 11 genomes of *Candidatus* Jettenia.
- **Fig. S4.** The potential for carbohydrate metabolism of 30 cMAGs. Glycoside hydrolases (GHs); Glycosyl transferases (GTs); Polysaccharide lyases (PLs); Auxiliary activities (AAs); Carbohydrate esterases (CEs); Carbohydrate-binding modules (CBMs).
- **Fig. S5.** Comparison of the potential for nitrogen metabolism of 3 *Candidatus* Jettenia species of this study and other 26 members of anammox bacteria**.**
- **Fig. S6.** Circular representation of cMAG.29. From outside to inside: *tnpA* on negative strand, *tnpA* on positive strand, genomic islands, coding sequences (CDSs) or RNAs on positive strand, CDSs or RNAs on negative strand, GC Content, and GC Skew.

**Table S1.** Composition of the culture medium.

| Chemical | Per liter contained |
|---|---|
| $KHCO_3$ | 1.0–2.5 g |
| $NaNO_2$ | 0.5–1.4 g |
| $NH_4Cl$ | 0.4–1.0 g |
| $CaCl_2$ | 0.05–0.15 g |
| $MgSO_4.7H_2O$ | 0.05–0.15 g |
| $KH_2PO_4$ | 0.025 g |
| $FeSO_4.7H_2O$ | 0.005 g |
| EDTA-2Na | 0.005 g |
| Trace element solution | 1–1.25 mL |

**Table S2.** Composition of trace element solution.

| Chemical | Per liter contained (g) |
|---|---|
| EDTA-2Na | 19.100 |
| $ZnSO_4 \cdot 7H_2O$ | 0.430 |
| $CoCl_2 \cdot 6H_2O$ | 0.240 |
| $MnCl_2 \cdot 4H_2O$ | 0.990 |
| $CuSO_4 \cdot 5H_2O$ | 0.250 |
| $NaMoO_4 \cdot 2H_2O$ | 0.220 |
| $NiCl_2 \cdot 2H_2O$ | 0.190 |
| $Na_2WO_4 \cdot 2H_2O$ | 0.050 |
| $H_3BO_3$ | 0.014 |

**Table S3.** Assessment of the purity and quantity of genomic DNA.

| Feature | Value |
|---|---|
| DNA concentration (μg) | 74.0 |
| $OD_{260/280}$ | 2.12 |
| $OD_{260/230}$ | 2.00 |
| Total genomic DNA (μg) | 40.7 |

**Table S4.** Basic information of HiFi reads.

| Feature | Value |
|---|---|
| Total reads (million) | 3.2 |
| Total bases (Gb) | 26.7 |
| Largest length (bp) | 31,071 |
| N50 length (bp) | 8,908 |
| N90 length (bp) | 5,995 |
| Average length (bp) | 8,462 |
| Q20% | 100 |
| Average quality | Q29.5 |

**Table S5.** Basic information of Illumina reads.

| Feature | Value |
|---|---|
| Total reads (million) | 235.5 |
| Total bases (Gb) | 35.3 |
| Q20% | 98.95 |
| Q30% | 96.07 |

**Table S6.** The quality assessment of HiFi reads assemblies.

| Assembly method | Contig number | Contig length (bp) | N50 (bp) | N90 (bp) | Largest length (bp) | Minimum length (bp) |
|---|---|---|---|---|---|---|
| hifiasm-meta | 25,771 | 1,247,998,565 | 76,563 | 19,755 | 7,419,406 | 4,411 |
| Flye | 25,916 | 952,651,823 | 79,796 | 14,422 | 7,468,635 | 1,002 |
| Canu | 24,013 | 937,638,827 | 57,470 | 15,810 | 5,853,242 | 3,903 |

**Table S7.** Reference genomes of 26 anammox bacteria.

| NCBI accession number | Species name | Assembly level | Completeness (%)# | Contamination (%)# |
|---|---|---|---|---|
| GCA_021777055.1 | *Candidatus* Anammoxibacter sp. OFTM134 | Scaffold | 95.1 | 3.4 |
| GCA_021777075.1 | *Candidatus* Anammoxibacter sp. OFTM301 | Scaffold | 82.8 | 9.7 |
| GCA_021777105.1 | *Candidatus* Anammoxibacter sp. OFTM214 | Scaffold | 93.9 | 5.7 |
| GCA_012515235.1 | *Candidatus* Anammoximicrobium sp. AS06rmzACSIP_251 | Contig | 98.2 | 5.8 |
| GCA_020724075.1 | *Candidatus* Anammoxoglobus sp. DR5_60_7 | Scaffold | 73.9 | 3.6 |
| GCA_017347445.1 | *Candidatus* Brocadia pituitae | Complete Genome | 98.9 | 2.8 |
| GCA_000949635.1 | *Candidatus* Brocadia sinica | Contig | 97.8 | 1.7 |
| GCA_001753675.2 | *Candidatus* Brocadia sapporoensis | Scaffold | 92.2 | 0.6 |
| GCA_000296795.1 | *Candidatus* Jettenia caeni | Contig | 100.0 | 3.9 |
| GCA_013360885.1 | *Candidatus* Jettenia caeni | Contig | 96.7 | 3.3 |
| GCA_016861385.1 | *Candidatus* Jettenia caeni | Contig | 84.6 | 1.1 |
| GCA_022180965.1 | *Candidatus* Jettenia caeni | Contig | 100.0 | 3.3 |
| GCA_021650895.1 | *Candidatus* Jettenia sp. AM49 | Complete Genome | 95.6 | 1.7 |
| GCA_900696655.1 | *Candidatus* Jettenia sp. AMX2 | Contig | 98.9 | 2.8 |
| GCA_008363445.1 | *Candidatus* Jettenia sp. AMX1 | Contig | 95.6 | 3.3 |
| GCA_014338295.1 | *Candidatus* Jettenia sp. AMX1 | Scaffold | 100.0 | 3.9 |
| GCA_021405185.1 | *Candidatus* Jettenia sp. AMX1 | Scaffold | 95.6 | 2.8 |
| GCA_024434255.1 | *Candidatus* Jettenia sp. AMX1 | Scaffold | 93.4 | 2.8 |
| GCA_005524015.1 | *Candidatus* Jettenia ecosi | Contig | 100.0 | 3.5 |

| | | | | |
|---|---|---|---|---|
| CT030148, CT573071, CT573072, CT573073, CT573074* | First genome *Candidatus* Kuenenia stuttgartiensis | Contig | 97.8 | 0.6 |
| GCA_011066545.1 | *Candidatus* Kuenenia stuttgartiensis | Complete Genome | 97.8 | 1.7 |
| GCA_900232105.1 | *Candidatus* Kuenenia stuttgartiensis | Complete Genome | 97.8 | 1.7 |
| GCA_021646405.1 | *Candidatus* Loosdrechtia aerotolerans | Contig | 96.7 | 3.3 |
| GCA_000786775.1 | *Candidatus* Scalindua brodae | Contig | 92.7 | 2.3 |
| GCA_002443295.1 | *Candidatus* Scalindua japonica | Contig | 95.5 | 3.4 |
| GCA_002632345.1 | *Candidatus* Scalindua rubra | Contig | 90.5 | 2.3 |

* EMBL-EBI accession number

### Calculated by CheckM

**Table S8.** The prediction results by PlasClass.

| ID | Predicted value |
|---|---|
| cMAG.1 | 8.25E-03 |
| cMAG.2 | 5.05E-06 |
| cMAG.3 | 2.37E-05 |
| cMAG.4 | 6.83E-06 |
| cMAG.5 | 3.92E-07 |
| cMAG.6 | 1.56E-08 |
| cMAG.7 | 7.58E-09 |
| cMAG.8 | 1.48E-03 |
| cMAG.9 | 7.98E-06 |
| cMAG.10 | 4.05E-02 |
| cMAG.11 | 4.02E-07 |
| cMAG.12 | 5.95E-07 |
| cMAG.13 | 1.51E-06 |
| cMAG.14 | 1.01E-06 |
| cMAG.15 | 1.73E-05 |
| cMAG.16 | 2.05E-04 |
| cMAG.17 | 6.07E-02 |
| cMAG.18 | 5.69E-03 |
| cMAG.19 | 2.92E-05 |
| cMAG.20 | 3.52E-11 |
| cMAG.21 | 9.00E-08 |
| cMAG.22 | 7.65E-06 |
| cMAG.23 | 3.47E-12 |
| cMAG.24 | 2.65E-09 |
| cMAG.25 | 4.30E-15 |
| cMAG.26 | 2.24E-09 |
| cMAG.27 | 3.43E-02 |
| cMAG.28 | 6.64E-04 |
| cMAG.29 | 3.22E-04 |
| cMAG.30 | 1.33E-03 |

### The threshold is 0.5.

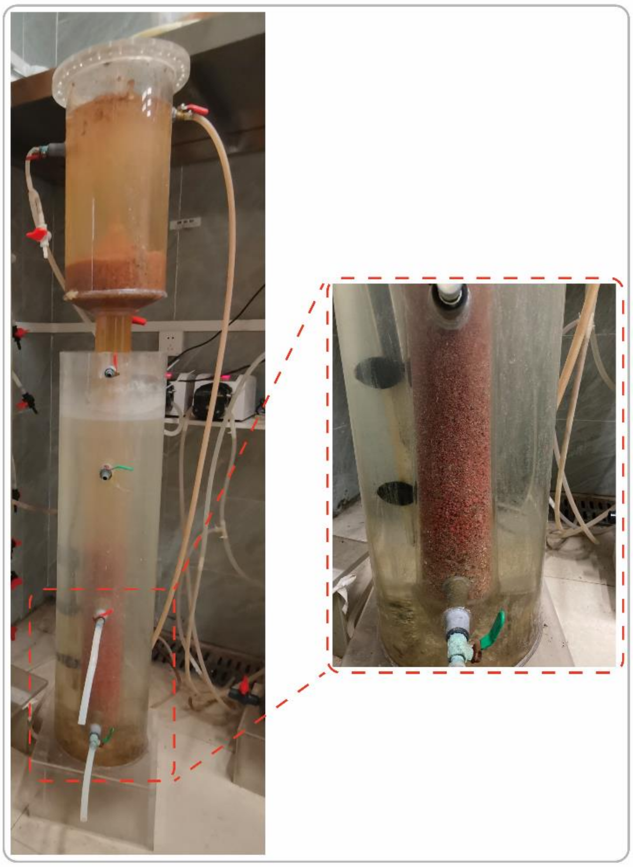

**Fig. S1.** The lab-scale expanded granular sludge blanket (EGSB) reactor.

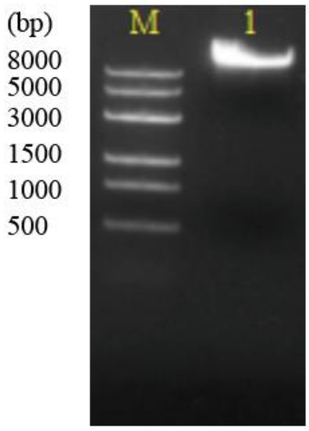


**Fig. S2.** Agarose gel electrophoresis of qualified genomic DNA.

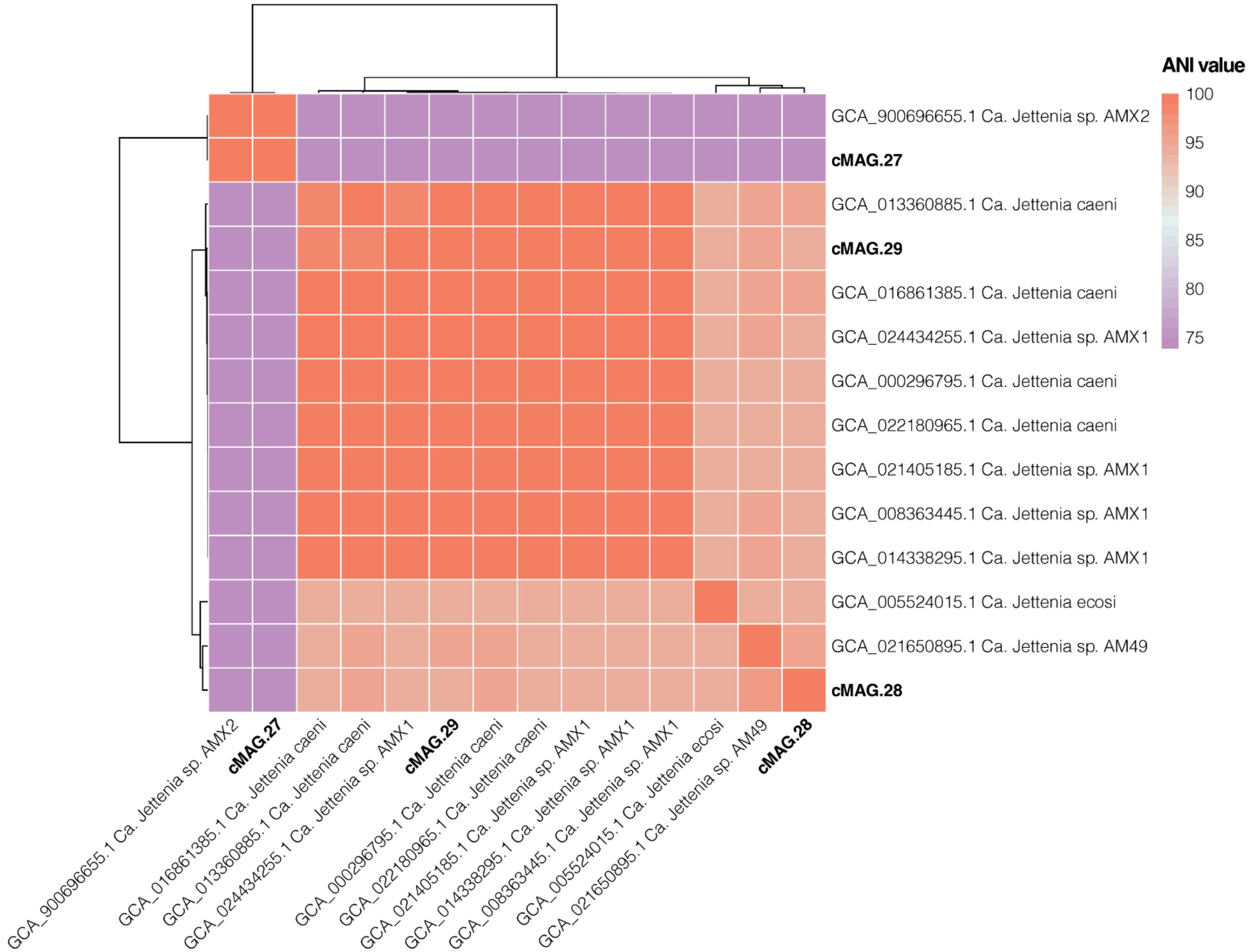


**Fig. S3.** The average nucleotide identity (ANI) value heatmap among 3 cMAGs and other 11 genomes of *Candidatus* Jettenia.

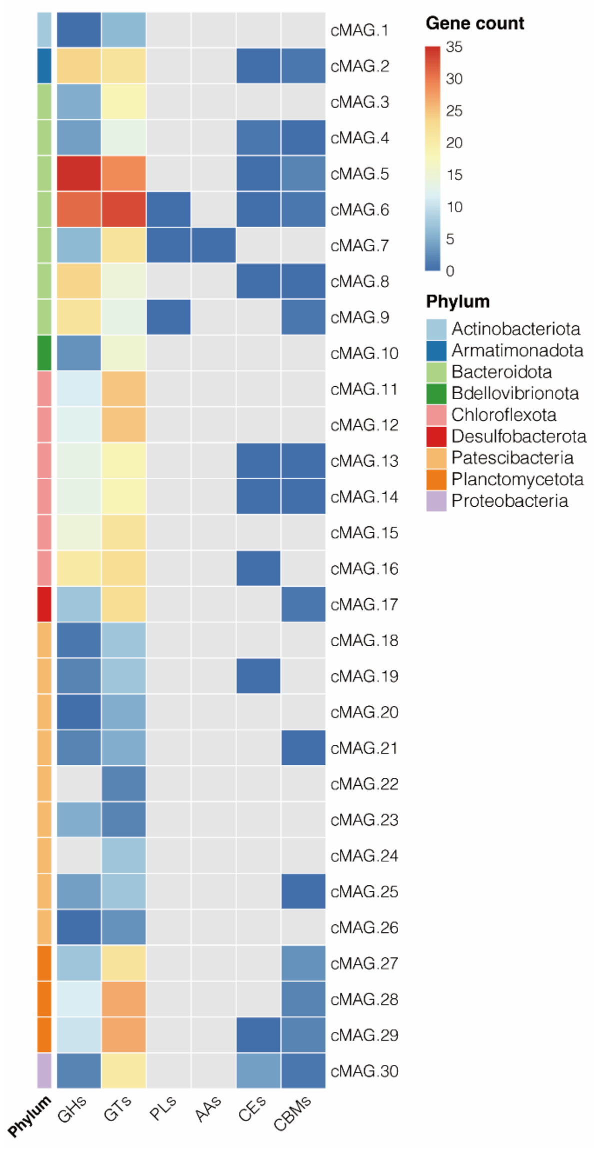


**Fig. S4.** The potential for carbohydrate metabolism of 30 cMAGs. Glycoside hydrolases (GHs); Glycosyl transferases (GTs); Polysaccharide lyases (PLs); Auxiliary activities (AAs); Carbohydrate esterases (CEs); Carbohydrate-binding modules (CBMs).

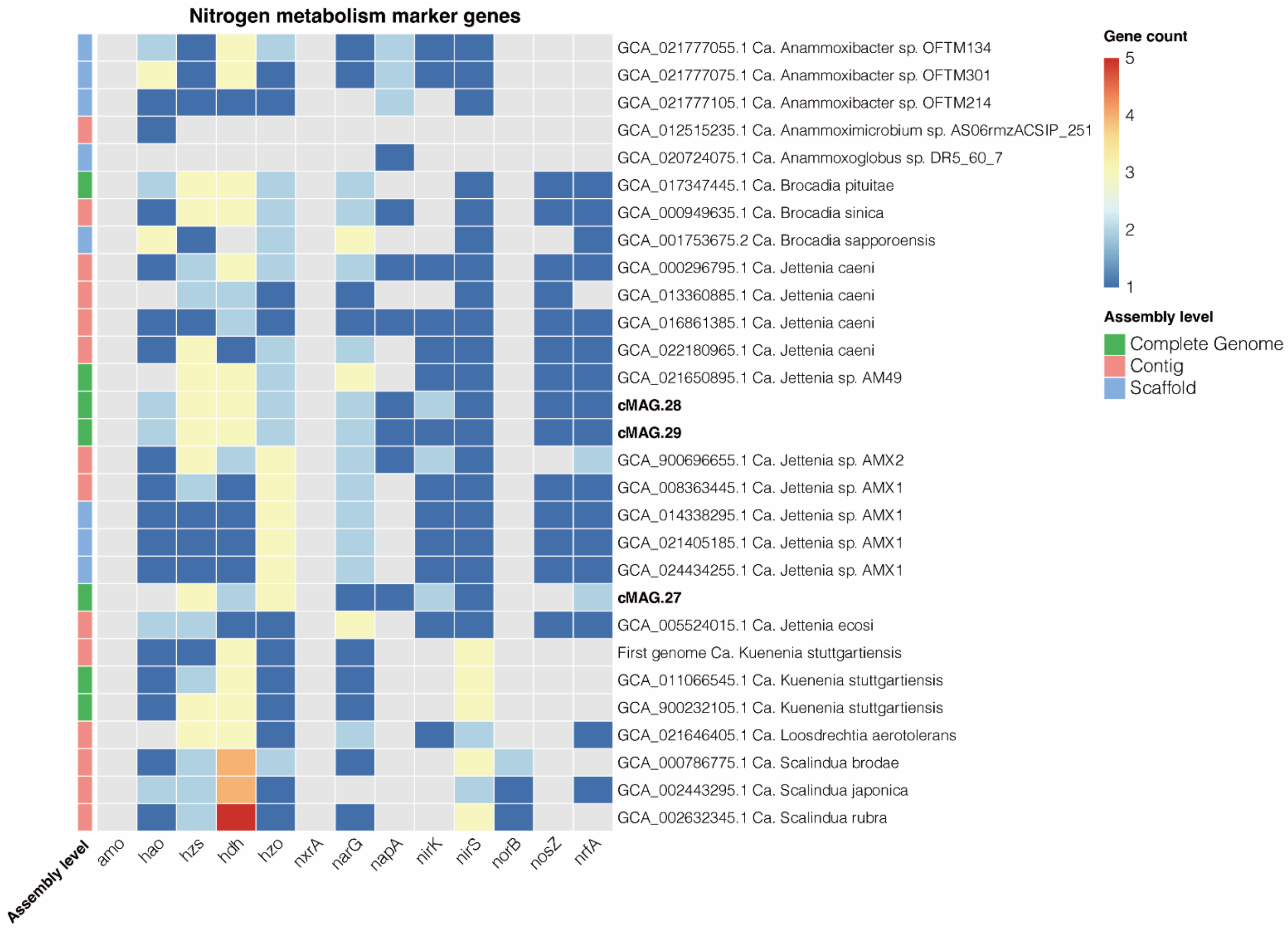


**Fig. S5.** Comparison of the potential for nitrogen metabolism of 3 *Candidatus* Jettenia species of this study and other 26 members of anammox bacteria.

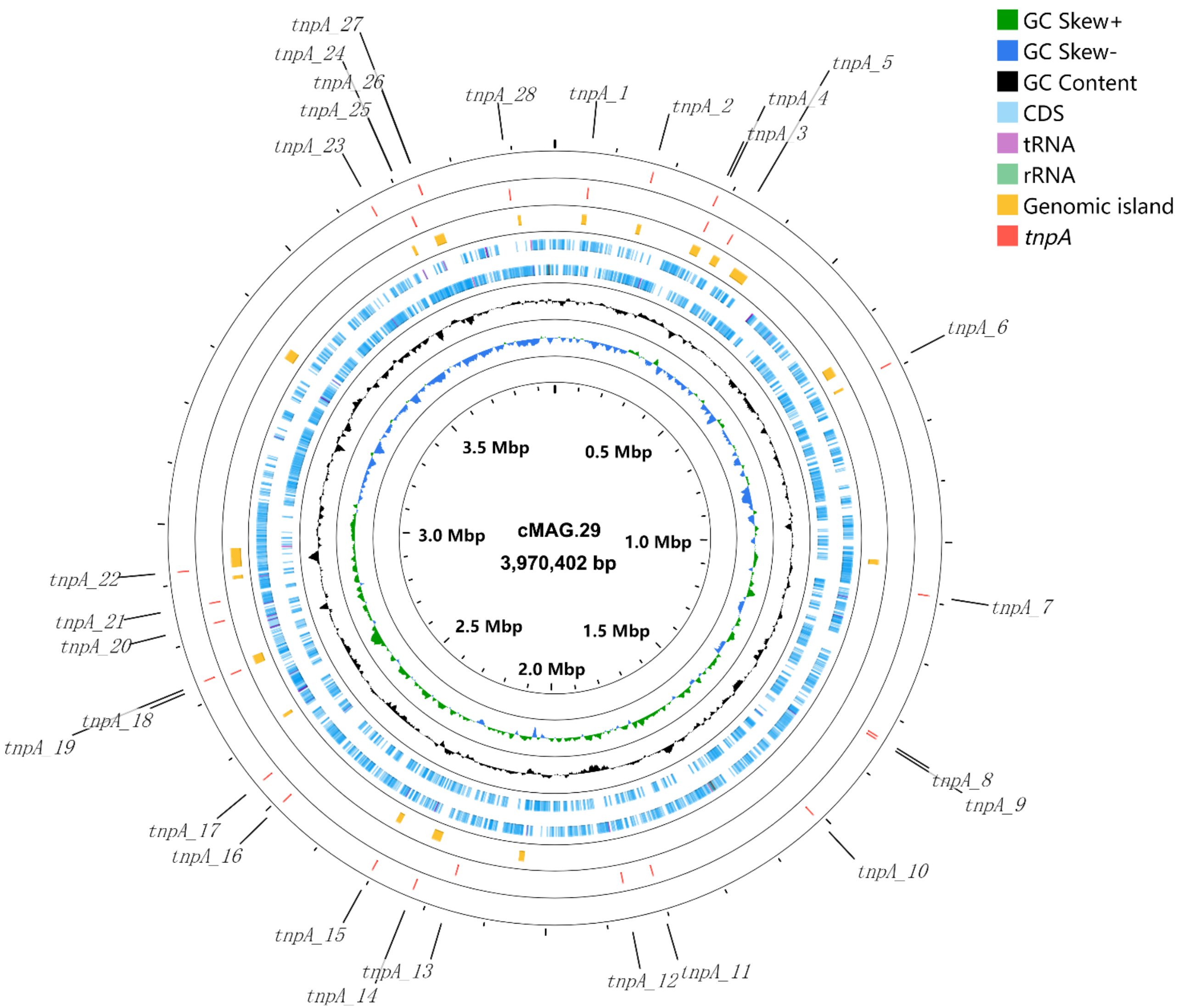


**Fig. S6.** Circular representation of cMAG.29. From outside to inside: *tnpA* on negative strand, *tnpA* on positive strand, genomic islands, coding sequences (CDSs) or RNAs on positive strand, CDSs or RNAs on negative strand, GC Content, and GC Skew.
